# Cotranslational folding and maturation of HIV-1 protease

**DOI:** 10.1101/2025.08.27.672612

**Authors:** Justin Westerfield, Felix Nicolaus, Ronald Swanstrom, Gunnar von Heijne

## Abstract

HIV-1 particle assembly depends critically on multiple proteolytic cleavages of viral polyproteins by the viral protease, PR. PR is translated as part of the Gag-Pro-Pol polyprotein, which undergoes autoproteolysis to liberate active, dimeric PR during virus particle maturation. Gag-Pro-Pol is produced via an infrequent -1 frameshifting event in ribosomes translating full length genomic RNA as Gag mRNA. Here, we study the cotranslational folding and autoproteolytic processing of frameshifted transframe-protease-reverse transcriptase (TF-PR-RT) constructs by *in vitro* translation. We demonstrate partial cotranslational folding of ribosome-bound PR at its conserved α-helix near the C terminus. Unexpectedly, we find that the initial dimerization of TF-PR-RT involves ribosome-bound nascent chains that are then not further cleaved. Moreover, only ribosome-bound nascent chains are substrates for PR-catalyzed processing. These observations are consistent with a model for virion assembly in which dimerization of a subset of Pro-Pol precursors leads to cleavage of PR monomers that then carry out the bulk of the proteolytic processing needed for virion maturation and infectivity.

## Introduction

Most retroviruses, including human immunodeficiency virus type 1 (HIV-1), assemble their virions at the plasma membrane of the infected cell ^1^. The precursor proteins Gag and Gag-Pro-Pol oligomerize on the inner face of the plasma membrane to nucleate budding of the enveloped particle ^2^. The Gag-Pro-Pol precursor is formed via a -1 frameshifting event that replaces the C-terminal domain of Gag with the transframe (TF) region in the -1 reading frame which allows translation to continue to include the Pro and Pol coding domains. The frameshift, however, is infrequent in that the Gag precursor is present in a 20-fold excess over the Gag-Pro-Pol precursor ^3,4^.

Virus particle production occurs with the Gag and Gag-Pro-Pol precursors (or even with Gag alone) but such particles are not infectious until further maturation. Both precursor proteins must undergo proteolytic processing by the encoded viral protease (PR, representing the Pro portion of Gag-Pro-Pol) to produce the mature proteins that function in formation of an infectious virion, along with other viral enzymes that function in the next infected cell ^5^. The Gag precursor encodes the structural proteins of the virus particle as well as the "late domain" that interacts with ESCRT proteins to effect budding of the host membrane surrounding the viral proteins ^6,7^, whereas the Gag-Pro-Pol precursor does not include the late domain but encodes the viral enzymes. Proteolytic processing of both precursors is an essential step in the virus life cycle, such that antiviral protease inhibitors can effectively inhibit HIV-1 maturation ^8,9^. PR, which is responsible for these processing events, is an aspartic proteinase and obligate homodimer ^5,10^. Thus, initial activation of protease activity requires dimerization of the Gag-Pro-Pol precursor, a process that may be enhanced by dimerization of the reverse transcription domain in the precursor ^11^.

There is evidence that Gag precursors can form low-order oligomers in the cytoplasm before moving to the plasma membrane where they oligomerize to induce budding ^12–20^. If Gag-Pro-Pol also participates in these low-order oligomers through their Gag domains, this would limit the chance of forming Gag-Pro-Pol homodimers in favor of Gag/Gag-Pro-Pol heterodimers, thus precluding PR dimerization and activation prior to membrane-associated oligomerization and budding. The rescue of a membrane-targeting mutant of Gag-Pro-Pol by wild type Gag is consistent with an interaction prior to reaching the membrane ^21,22^. The absence of protease activation from dimerization in the cytoplasm can be inferred by the observation that if a head-to-tail PR dimer is encoded in the Gag-Pro-Pol precursor, it efficiently initiates premature proteolytic processing ^23^. In addition, when a tight-binding, non-nucleoside reverse transcriptase inhibitor (NNRTI) is used to potentiate dimerization of the reverse transcriptase domain (RT; encoded in the Pol portion of Gag-Pro-Pol) this also prematurely activates the viral protease ^24–28^. This NNRTI-induced premature dimerization appears to happen only after the Gag-Pro-Pol precursor has migrated to the membrane where Gag and Gag-Pro-Pol engage in higher order oligomerization ^29^. When premature activation does occur, it leads to a nonproductive pathway for virion formation.

The protease becomes active in the context of a dimerized Gag-Pro-Pol precursor. The Pro-Pol domain is released from the upstream portion of Gag by a cleavage at the same site that is first cleaved in Gag (NC/SP2) ^30,31^. This cleavage event is followed by cleavage of a site within the transframe region upstream of the PR domain ^30,32–36^; importantly, evidence from an *in vitro* translation system indicates that these two upstream cleavage events occur in *cis* by the embedded dimerized protease ^31^.

Since the protease can exist in a monomer form (initially in the Gag-Pro-Pol precursor) and as a dimer, this has led to studies of protease structure as a monomer ^37–39^. Also, while the Pro-Pol domains can dimerize in their precursor form, how this structure proceeds to give mature protease (the dimer of identical 99 amino acid subunits) that can then carry out cleavage at five sites in Gag and four sites in Pro-Pol is an active area of research. The transframe (TF) domain, the -1 reading frame of Gag present upstream of Pro, has been suggested to inhibit protease activity until it is cleaved from the N terminus of the protease ^35,40,41^. One line of evidence has described flexibility within the PR structure that allows the N-terminal protease cleavage site to be cleaved in *cis* followed by cleavage of the C terminal protease cleavage site as the mechanism of generation of mature protease from the precursor ^41–45^. The released mature PR dimer is similar in catalytic activity to model precursor forms of PR with short ^41^ but not long extension ^42^ beyond the protease cleavage sites, but even short extensions destabilize the protease dimer ^41^, suggesting that it is the mature protease that is responsible for most of the catalytic activity in the virion. Despite these lines of evidence, dimerization-induced activation of PR within the Gag-Pro-Pol precursor and the potential cotranslational folding of PR itself is a process that is incompletely understood.

In this report we have used Force Profile Analysis (FPA) in an *in vitro* translation system to examine cotranslational protein folding of PR, measured as the ability of the folding reaction to exert a pulling force on a downstream translational arrest peptide ^46^. We find that the PR domain embedded in a truncated TF-PR-RT context has little structure until the C-terminal α-helix emerges from the ribosome exit tunnel. The formation of the α-helix does not appear to nucleate upstream folding, as the folding force was unchanged when mutations to the hydrophobic core of PR were made. Unexpectedly, we further find that only nascent, ribosome-bound TF-PR-RT chains can, first, initiate productive PR dimerization, and second, be cleaved by PR, while released, presumably dimerized, chains are largely refractory to proteolysis. Applying these findings to the situation in the infected cell, we propose a model for PR activation where infrequent precursor dimerization events during budding generate TF-Pro-Pol dimers not bound to the plasma membrane and this leads to cleavage of PR monomers *in trans* that then dimerize to become the highly active form for virion maturation.

## Materials and methods

### Enzymes and chemicals

Enzymes and other reagents were purchased from Thermo Fisher Scientific (Waltham, Massachusetts, U.S.), New England BioLabs (Ipswich, Massachusetts, U.S.), and Sigma-Aldrich (subsidiary of Merck Life Science, Darmstadt, Germany). Oligonucleotides were ordered from Eurofins Genomics (Luxembourg City, Luxembourg) and Thermo Fisher Scientific. L-[^35^S]-methionine (NEG009T001MC) was provided by PerkinElmer (Shelton, Connecticut, U.S.).

### Cloning and Mutagenesis

The DNA sequence of Gag-Pro-Pol was acquired from the pNL4-3 clone of HIV-1 ^47^. To focus on PR and its immediate fusion neighbors, we used a synthetic gene that begins with TF already in the -1 frame and ends at RT residue 30. A variable-length linker, the 17-residue SecM(*Ec*) arrest peptide, and a 23-residue C-terminal tail were added to the C-terminus, Fig. 1a, and the resulting ORF was subcloned downstream of a T7 promoter in a pET-19b vector carrying an ampicillin resistance cassette. Simple mutagenesis was performed by mismatch PCR, whereas more extensive mutagenesis was performed with a combination of PCR and Gibson assembly ^48^. See Supplemental Table S1 for amino acid sequences of all constructs.

**Figure 1.**
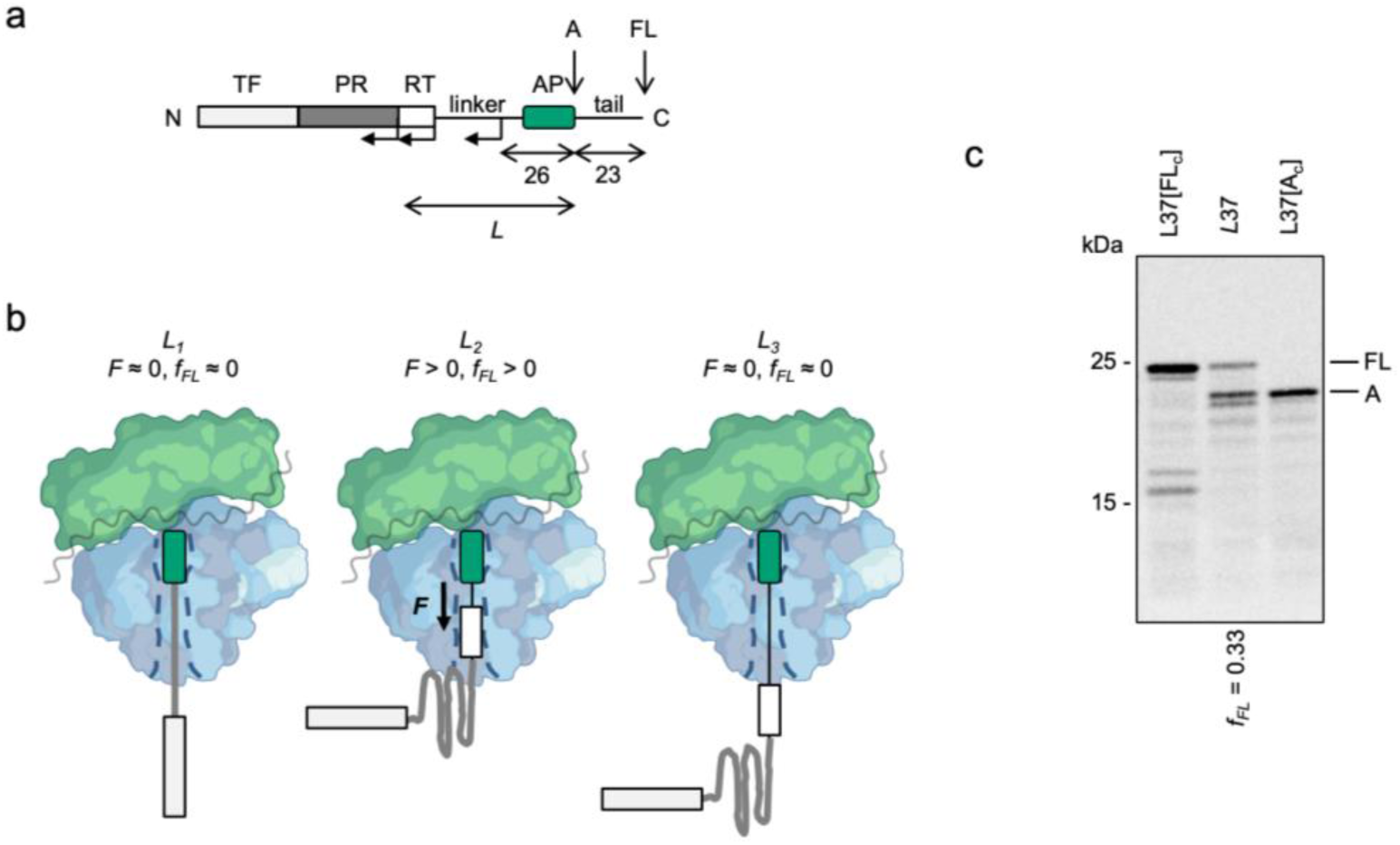
The force profile assay. (a) Basic construct (*L* = 60 residues). The HIV-1 transframe (TF), protease (PR), reverse transcriptase residues 1-30 (RT), linker, SecM(*Ec*) AP, and C-terminal tail are indicated, together with the arrested (*A*) and full-length (*FL*) products. *L* denotes the number of residues between the C-terminal end of PR and the last residue in the AP. The 23-residue tail and a 26-residue segment including the AP have the same sequence in all constructs. Shorter constructs were made by progressively deleting residues in, first, the RT, then the linker, and last the C-terminal PR regions (angled arrows), as detailed in Supplementary Fig. S1. (b) At construct length *L_1_*, PR (gray) is located too deep in the exit tunnel to be able to fold, no pulling force *F* is generated, and little full-length product is produced (*F* ≈ 0, *f_FL_* ≈ 0). At *L_2_*, PR is starting to fold, generating a strong pulling force and more full-length product (*F* > 0, *f_FL_* > 1). At *L_3_*, PR has been fully folded and little pulling force is generated (*F* ≈ 0, *f_FL_* ≈ 0). (c) SDS-PAGE gel showing *A* and *FL* products for the *in vitro* translated, [^35^S]-Met labelled construct with *L* = 37 residues. Control constructs L37[A_c_] and L37[FLc] have, respectively, a stop codon and an inactivating Ala codon replacing the last Pro codon in the AP, leading to the production of exclusively *A*-sized and *FL*-sized proteins. Mw markers are indicated on the left.

### In vitro pulse-labeling analysis

Before expression, linear DNA spanning the open reading frame from constructs was generated by PCR using primers against the T7 promoter (5’-CCCGCGAAATTAATACGACTCACTATAGGG-3’) and terminator (5’-GCTAGTTATTGCTCAGCGG-3’), respectively, followed by PCR clean-up using a Thermo Scientific (Waltham, Massachusetts, U.S.) GeneJET PCR purification kit (catalog number K0701). The linearized DNA was then expressed using the New England BioLabs (Ipswich, Massachusetts, U.S.) PURExpress *in vitro* protein synthesis kit (catalog number E6800L). Briefly, 2.2 µL of linearized DNA was mixed with 0.8 µL [^35^S]-Met, 4 µL solution A and 3 µL solution B, and the mixture was incubated at 37°C for 15 min shaking at 750 RPM in a thermomixer. To stop the reaction and precipitate the proteins, 10 µL of 10% trichloroacetic acid was added and the mixture was placed on ice for 30 min. The precipitate was centrifuged at >20,000 x g for 10 min, and then the supernatant was aspirated off. To resuspend the proteins, 12 µL of SDS-sample buffer was added and the reaction tube was shaken for 10 min at 37°C.

For time-course experiments, reactions were carried out in the same way as above with a few differences. The purified, linearized DNA was adjusted such that the final reaction concentration was 10 ng/µL, as this was found to affect cleavage time. Also, a much larger reaction mixture was made (total 40 µL), from which 5 µL samples were taken at denoted time points and transferred to tubes containing 5 µL of 10% trichloroacetic acid. After the first sample was removed, excess non-radioactive Met was added to a final concentration of 10 mM. Further treatments were the same for all samples.

The PR inhibitor darunavir (Sigma-Aldrich SMLO937) was dissolved at 18 mM in DMSO, and, 0.5 µL was added at the beginning of the PURE reaction where applicable.

For experiments in the presence of purified PR, PURE *in vitro* translation reactions were carried out as above, but with 1 µL of diluted, purified PR added either at the start of the reaction or at the start of the chase, as denoted in the figures. Purified PR (a gift from Dr. Celia Schiffer) was diluted into *in vitro* translation buffer (9 mM Mg(OAc)_2_, 5 mM K-phosphate pH 7.3, 95 mM K-glutamate, 5 mM NH_4_Cl, 0.5 mM CaCl_2_, 1 mM spermidine, 8 mM putrescine, 1 mM DTT) such that the final concentration in each reaction was 1 µM.

Samples (for both single reactions and time-course experiments) were then incubated with 0.25 mg/ml RNase for 30 min. at 37°C to hydrolyze tRNA, and the proteins were subsequently separated by SDS-PAGE. Gels were fixed in 30% (v/v) methanol or ethanol and 10% (v/v) acetic acid. After incubation in Gel-Dry^TM^ Drying Solution (Invitrogen, Thermo Fisher Scientific, Waltham Massachusetts, U.S.) for 30 min, gels were dried in a Hoefer GD 2000 gel dryer (Hoefer Inc., U.S.). After exposing dried gels to phosphorimaging plates (BAS-IP, Fujifilm, Tokyo, Japan) overnight, plates were scanned using a Fuji FLA-9000 imager (Fujifilm, Tokyo, Japan) to collect radio-images of gels. Band intensity profiles were obtained using the Fiji (ImageJ) software ^49^ and quantified with the in-house software EasyQuant (https://github.com/gvh-lab/easyquant) in order to determine the fraction full-length protein *f_FL_* = *I_FL_*/(*I_FL_* + *I_A_*), where *I_A_*, *I_FL_* are the intensities of the *A* and *FL* bands, respectively, Fig. 1c. A_c_ and/or FL_c_ size controls were included in the SDS-PAGE to confirm the identity of the *A* and *FL* bands. Each data point in the force profile represents the average of at least three independent replicates (*i.e*., independent PURE *in vitro* translation reactions) and includes the standard error of the mean (SEM).

Molecular weights were calculated using the Expasy Compute pI/Mw tool at https://www.expasy.org.

To extract lag-time midpoints from pulse-chase experiments, the fraction of cleaved protein *f_cleaved_ = I_PR_ /(I_PR_+ I_A_*) was first determined, where *I_PR_* is the integrated intensity of the released PR, and as above, *I_A_* is the integrated intensity of the *A* band, Fig. S1. The resulting *f_cleaved_* as a function of chase time was fit to a simple two-state sigmoidal curve:

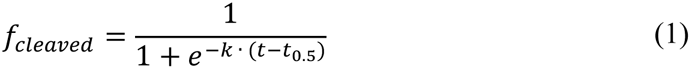

where *k* is the slope of the transition and *t_0.5_* is the lag-time midpoint Supplemental Fig. S2.

To determine arrest-peptide release rates (*k_R_*) and release-time midpoints (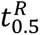), the fraction arrested *f_A_*(*t*) (= 1–*f_FL_*(*t*)) was fit to a single exponential decay function,

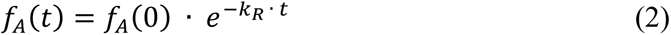

where *f_A_(t)* is the fraction of arrested protein at chase time *t*, and *f_A_(0)* is the fraction of arrested at time zero (*i.e*., at the beginning of the chase). Pulse-chase curves were fit using homemade Python scripts invoking SciPy ^50^ non-linear least squares fitting. Arrest peptide release-time midpoints were calculated as 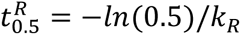.

## Results

### Force Profile Analysis

FPA takes advantage of the fact that translational arrest peptides (APs) can induce a temporary pause in translation by binding tightly to the upper part of the ribosome exit tunnel ^51,52^. AP-induced ribosome stalling can be overcome by pulling forces acting on the nascent chain ^53,54^, with different APs being sensitive to different force levels ^55^. Such pulling forces can be generated by, *e.g*., cotranslational protein folding or cotranslational insertion of transmembrane segments into a membrane ^46,56,57^. Therefore, APs can be conveniently used as force sensors to study cotranslational events *in vitro* and *in vivo*.

Here, we have used a series of plasmid constructs to produce increasingly longer portions of the frameshifted HIV-1 TF-PR-RT precursor protein, with the C-terminal end of PR placed *L* residues upstream from the C-terminal end of the *E.coli* SecM (SecM(*Ec*)) AP, which in turn is followed by a 23-residue C-terminal tail ^58^, Fig. 1a (see Supplemental Table S1 for amino acid sequences of all constructs). By varying *L*, PR can be positioned in different locations in the ribosome exit tunnel at the point when the ribosome reaches the last codon of the AP, and this will therefore generate pulling forces of different magnitude depending on whether the peptide chain can fold or not in that location. In a construct where PR can start to fold at the point when the ribosome reaches the last codon of the AP, Fig. 1b (middle), a strong pulling force *F* will be exerted on the nascent chain, which in turn reduces translational pausing at the AP and results in mostly full-length protein (including the C-terminal tail) being produced during a short pulse with [^35^S]-Met. In contrast, in constructs where PR is located too far up the exit tunnel to be able to fold, or has already folded before the point when the ribosome reaches the end of the AP, little force is exerted on the AP, translational pausing will be efficient, and more of the arrested form of the protein will be produced during a short pulse with [^35^S]-Met, Fig. 1b (left and right). After *in vitro* translation in the *E. coli*-derived PURE coupled transcription-translation system ^59^ for 15 min., full-length (*FL*) and arrested (*A*) protein species are separated on a SDS-PAGE gel and the fraction full-length protein is calculated as *f_FL_* = *I_FL_*/(*I_FL_*+*I_A_*), where *I_FL_* and *I_A_* are the intensities of the bands representing *FL* and *A* species, respectively, Fig. 1c. Because *f_FL_* is sensitive to pulling force, it can be used as a proxy for *F* ^54,60–62^. A force profile (FP), in which *f_FL_* is plotted against *L* (*i.e.*, each construct generates one data point), thus can be used to follow the cotranslational folding of PR.

### PR undergoes a cotranslational folding event starting at L ≈ 25 residues

In order to detect cotranslational folding events in PR as it emerges from the ribosome exit tunnel, a FP was determined using a series of constructs where *L* was varied between 18 and 60 residues (named L18 to L60), Fig. 2a (see Supplemental Table S1 for sequences); for some *L* values we also made control constructs in which the AP was inactivated either by mutating its C-terminal Pro to Ala (FL_c_ controls, producing only full-length protein), or by mutating its C-terminal Pro codon to a stop codon (A_c_ controls, producing only arrested-size protein), c.f., Fig. 1c. Note that constructs with *L* ≤ 24 residues have short C-terminal deletions in PR, Supplemental Table S1.

**Figure 2.**
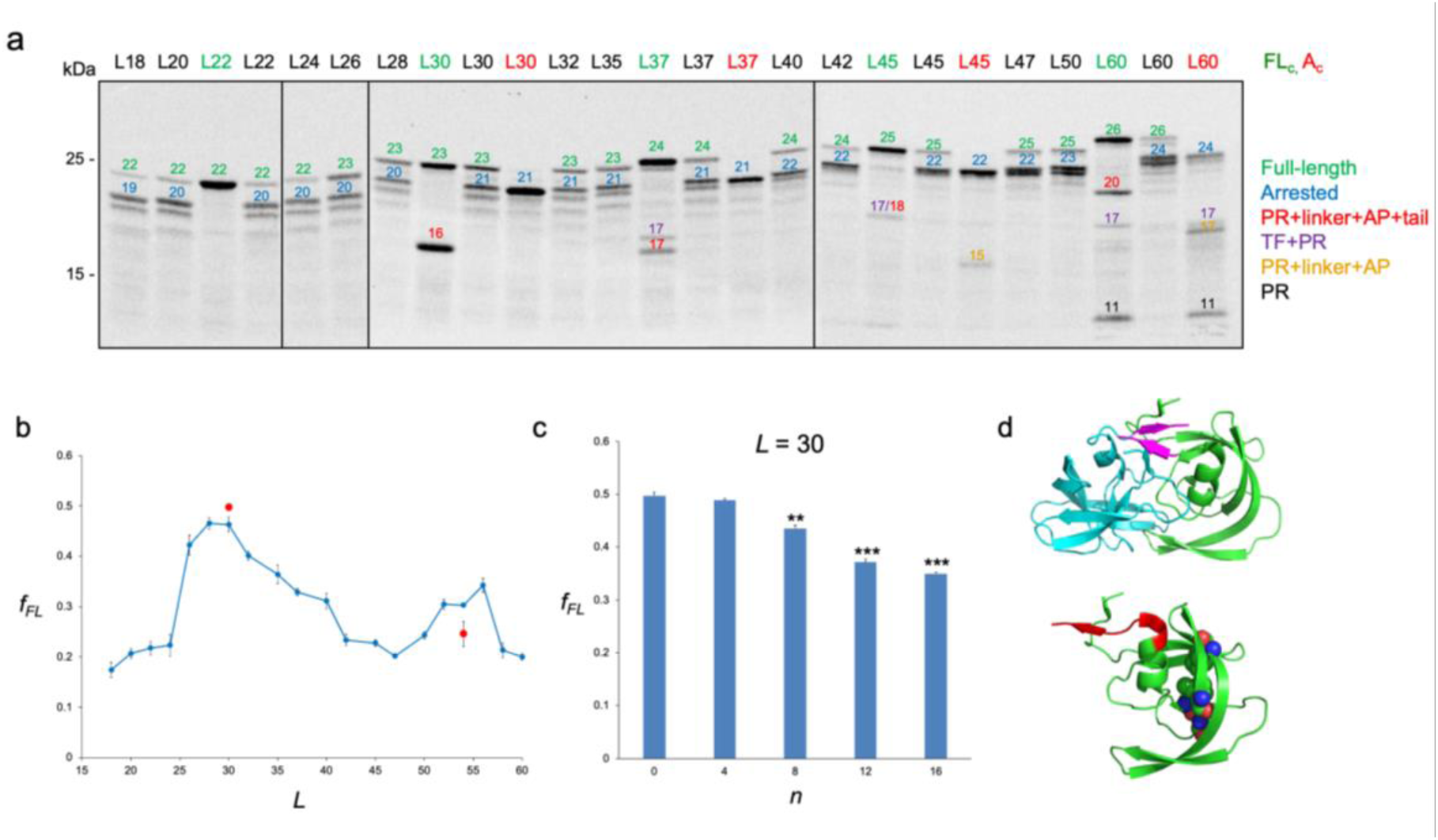
PR force profile. (a) 15 min. *in vitro* translations of constructs with an active AP (constructs indicated in black), FL_c_ controls (constructs indicated in green), and A_c_ controls (constructs indicated in red). Full-length, arrested, and proteolysis products are indicated by their calculated Mw’s (color-coded as indicated on the right). (b) FP for constructs with *L* = 18 to 60 residues (See Supplemental Table S1 for sequences). The red data points are for the quadruple PR mutant L^33^D-I^64^D-V^75^D-V^77^D. Each data point represents the average ± SEM of at least three replicates (*i.e*., independent PURE *in vitro* translation reactions). (c) *f_FL_* values for the *L* = 30 construct with increasing numbers, *n*, of C-terminal residues replaced by alternating GS repeats (e.g., *n* = 4 means GSGS). Residues replaced in the *n* = 8 mutant are indicated in red in *d*, lower panel. (d) Upper panel: 3D structure of the PR dimer (PDB ID 4QJ6). The C-terminal β-strand is shown in magenta. Lower panel: one PR monomer showing the eight C-terminal residues (in red) and the locations of the residues mutated in the quadruple PR mutant (in spacefill).

As seen in Fig. 2b, *f_FL_* increases sharply between *L* = 24 and *L* = 26 residues, is maximal at *L* = 28-30 residues, and returns to baseline at *L* = 42 residues, indicating a cotranslational folding event. A second peak of lower amplitude is seen for *L* = 50-56 residues.

To probe the nature of the folding events causing the two peaks, we mutated the L30 construct by replacing an increasing number of C-terminal PR residues with Gly-Ser (GS) repeats. This led to a significant decrease in *f_FL_* values when 8 or more residues were replaced, Fig. 2c. Replacement of the C-terminal β-strand (T^95^NLF^99^) with GSGS thus has no effect on *f_FL_* in the L30 construct, but extending the replacements into the C-terminal α-helix (residues 86-94) reduces *f_FL_*, Fig. 2d. We also simultaneously mutated four residues (L^33^, I^64^, V^75^, V^77^) in the hydrophobic core of the PR dimer structure to D in the L30 and L54 constructs, Fig. 2d. This had no effect on *f_FL_* for the L30 construct, but significantly reduced *f_FL_* for the L54 construct, Fig. 2b (red data points).

We conclude that the first peak in the FP is caused by a cotranslational folding event that initiates when the C-terminal end of PR reaches ∼25 residues from the PTC. This folding event does not seem to involve the entire PR monomer, however, but appears to be caused mainly by the formation of the C-terminal α-helix which at that point has its C-terminal end located ∼30 residues from the PTC, within the vestibule of the exit tunnel. The folded form of a monomeric PR mutant lacking the dimerization-mediating, short C-terminal β-strand (T^95^NLF^99^) is only marginally stable ^38,63^, and the monomer may be further destabilized when held in proximity to the ribosome ^64–66^, possibly explaining the absence of a folding event involving the whole PR domain.

The second peak, at *L* = 50-56 residues, corresponds to a situation where the entire PR domain is exposed outside the exit tunnel, as only ∼35-40 residues of an extended nascent chain are embedded within the ∼100Å long tunnel ^67,68^. At these chain lengths, it is possible that the PR segment on the arrested ribosome can reach and dimerize with the PR segment on the ribosome stacked behind it on the mRNA, or with an already synthesized and released TF-PR-RT chain ^69^, causing the observed increase in *f_FL_*. A dimerization event would be consistent with the observed effect on *f_FL_* of the hydrophobic core mutations in the L54 construct, although we cannot rule out other causes for this peak.

### In vitro-translated TF-PR-RT constructs produce active PR

As noted above, while measuring the FP we also expressed control constructs with mutant arrest peptides which do not arrest but are released normally at either full-length (FL_c_) or arrested length (A_c_). As seen in Figure 2a, various cleavage products were observed for the *L* ≥ 30 residues FL_c_ constructs and for the *L* ≥ 45 residues A_c_ constructs, suggesting the formation of active PR; notably, however, no such cleavage products were seen during the 15 min. translation reaction for any of the constructs with an active AP that are retained on the ribosome. These observations piqued our interest, and we decided to look more deeply into the processing of the TF-PR-RT constructs.

Constructs with *L* ≤ 24 residues do not contain the C-terminal β-strand (T^95^NLF^99^) required for dimerization of PR ^37^, and hence are not expected to produce active enzyme. Moreover, while all constructs contain the cleavage site SFSF/PQIT between TF and PR, only constructs with *L* ≥ 34 residues contain the native cleavage site TLNF/PISP between PR and RT (Supplemental Table S1). Consistent with this, for L30[FL_c_] we see only one cleavage product with a Mw corresponding to that expected for the PR+linker+AP+tail fragment (16.4 kDa; Fig. 2a), while for L37[FL_c_] we see two cleavage products of Mw of ∼17 kDa, close to the expected 17.2 kDa for the PR+linker+AP+tail fragment and 17.3 kDa for the TF+PR fragment (*c.f*., Fig. 1c). In addition, for L60[FL_c_] there is a band at ∼11 kDa which is approximately the Mw of mature PR (10.8 kDa). In contrast, there are no cleavage products seen for L30[A_c_] and L37[A_c_], but a fragment with the size expected for a PR+linker+AP fragment (15.5 kDa) is seen for L45[A_c_], and two bands at ∼17 kDa, close to the Mw’s expected for the TF+PR and PR+linker+AP fragments, and one band at the Mw expected for PR, are seen for L60[A_c_]. The expected TF fragment has a calculated Mw of 6.5 kDa and is too small to be visible on the gels.

To confirm that the observed fragments are produced by PR-mediated cleavage of the FL_c_ and A_c_ products, we repeated the experiments with the inactivating PR(D^25^N) mutation ^70^. As seen in Fig. 3a, no cleavage products were seen for the tested constructs in this case. Thus, the *in vitro* translation reactions produce active PR, at least for the FL_c_ and the longer A_c_ constructs.

**Figure 3.**
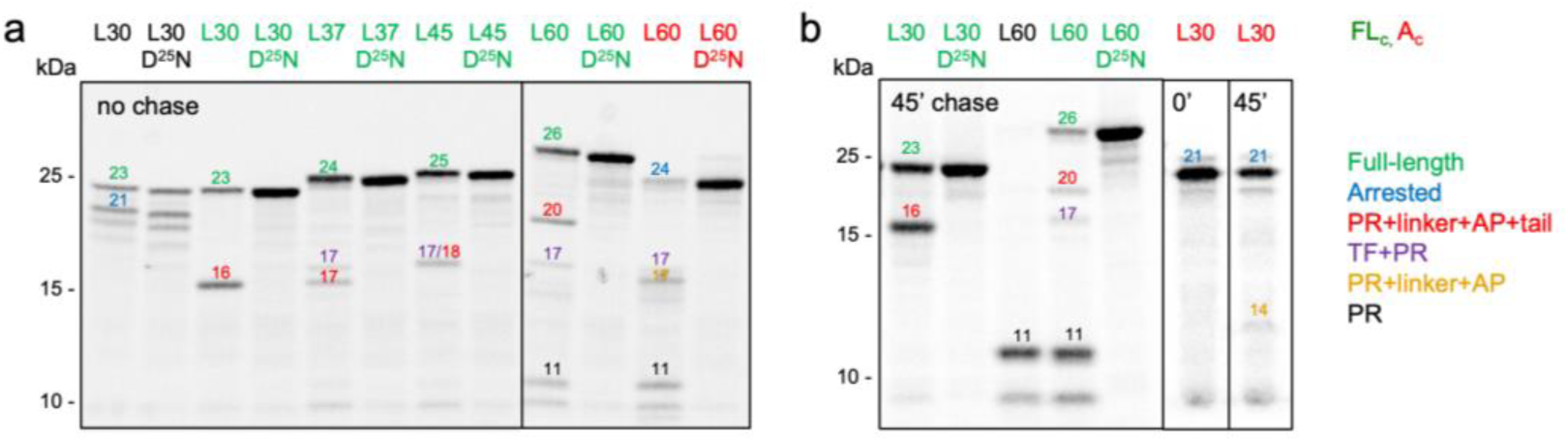
Autoproteolytic cleavage of TF-PR-RT constructs. (a) SDS-PAGE gels of *in vitro* translated, [^35^S]-Met labelled constructs (indicated in black), together with *A_c_* (indicated in red) and *FL_c_* (indicated in green) control constructs. Cleavage products are identified by their Mw’s according to the color code on the right. (b) Same as in panel *a*, but after a 45 min. chase with excess non-radioactive Met. The ∼9.5 kDa fragment at the bottom of the gels is a contamination from the PURE system itself, see Fig. 6a. The L30[A_c_] lanes in panel *b* are also shown in Fig. 6b.

### Ribosome-attached nascent TF-PR-RT chains are susceptible to cleavage by PR but released polyprotein is not

To analyze the cleavage reactions in more detail, we performed an initial set of pulse-chase experiments on L30[FL_c_], L30(D^25^N)[FL_c_], L30[A_c_], L60, L60[FL_c_], and L60(D^25^N)[FL_c_]. Non-radioactive Met was added in ∼15,000-fold excess after the standard 15 min. incubation with [^35^S]-Met, and samples were collected after an additional 45 min. chase. The relative amounts of cleavage products remained approximately the same for L30[FL_c_] and L60[FL_c_] before and after the chase, Fig. 3a, b. In marked contrast, L30[A_c_] was largely refractory to autoproteolysis, and only a small amount of processed PR+linker+AP was seen after the chase, Fig. 3b. Thus, L30[A_c_] produces only marginal amounts of catalytically active protease during the 45 min. chase, whereas the closely related L30[FL_c_] construct that includes the 23-residue C-terminal tail produces ample amounts even without a chase. The most obvious difference between these two constructs is that the entire TF-PR segment (located 53 residues from the stop codon in L30[FL_c_]) is exposed outside the ribosome exit tunnel for a time corresponding to the translation of ∼10-15 3’ codons (*i.e*., ∼10-20 sec ^71^) before synthesis of the L30[FL_c_] chain terminates and it is released from the ribosome, while the C-terminal β-strand in PR (located 29 residues from the end of L30[A_c_]) does not appear outside the exit tunnel at any point before chain termination of L30[A_c_]. This suggests that the initial TF-PR dimerization reaction – the first step in the production of active PR – involves at least one ribosome-bound nascent chain.

Strikingly, the ribosome-arrested L60 construct, which did not show any cleavage products after a 15 min. translation reaction (Fig. 2a), was efficiently converted to mature ∼11 kDa PR during the chase, Fig. 3b, unless the PR inhibitor darunavir ^72^ was included in the reaction, Supplemental Fig. S1. Furthermore, L60 undergoes complete autoproteolysis to PR, showing none of the intermediate cleavage products observed in the non-arrested L60[FL_c_] control, Fig. 3b. This suggests that ribosome-bound nascent chains are the main PR substrates, with released chains being largely refractory to proteolysis.

A more extensive pulse-chase analysis of a range of ribosome-arrested constructs (*L* = 30-60 residues), Fig. 4, showed typical autoproteolytic cleavage kinetics, with a slow initial lag phase during which the first, catalytically active PR molecules are produced (presumably via dimerization of the TF-PR-RT precursor and formation of the first highly active PR dimers), followed by rapid processing of the arrested (but not full-length) form, first to the ribosome-bound PR+linker+AP fragment, and then to fully cleaved ∼11 kDa PR (see Supplemental Fig. S2 for quantitative analysis). As expected from the absence of an intact PR/RT cleavage site in the L30 construct, no ∼11 kDa PR was produced from L30 during the 45 min. chase (Fig. 4a), although the arrested form slowly converts to a ∼14 kDa fragment (lag-time midpoint t_0.5_ = 39 min.), which may correspond to the 14.0 kDa PR+linker+AP fragment ^54^. Notably, autoproteolytic cleavages are seen only for the arrested (*A*), ribosome-bound forms of the proteins, not for the released full-length (*FL*) products.

**Figure 4.**
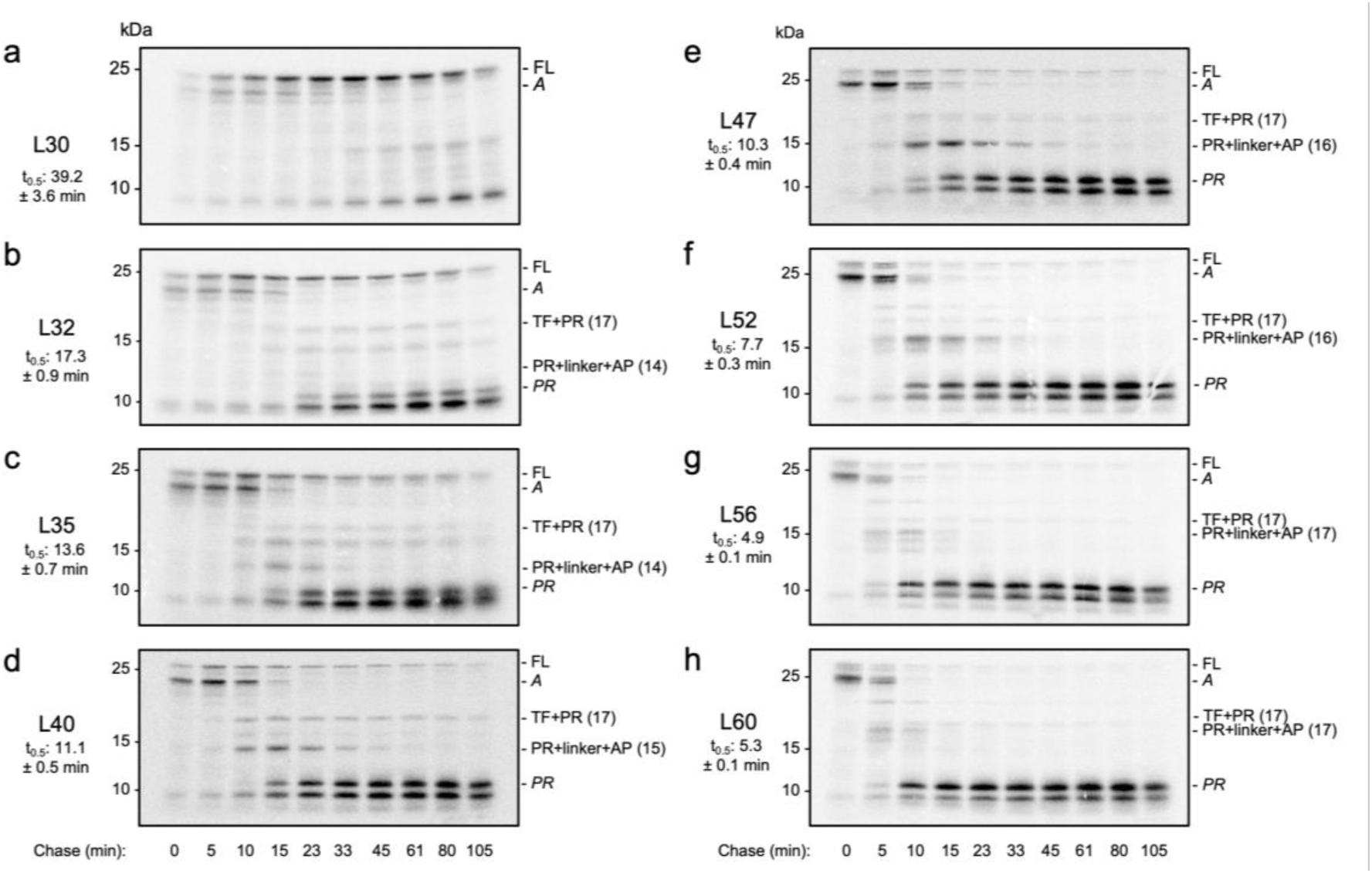
Pulse-chase analysis of the autoproteolytic cleavage reaction. (a) Constructs were translated *in vitro* for 15 min. in the presence of [^35^S]-Met and then chased after addition of excess non-radioactive Met for the indicated times. Full-length (*FL*), arrested (*A*), ∼11 kDa mature PR, TF+PR, and variable-length PR+linker+AP products are indicated (with calculated Mw’s). L30 is missing the PR/RT cleavage site, and therefore cannot generate mature PR. Lag-time midpoints (t_0.5_) ± the standard error of the fitting are indicated. See Methods and Supplemental Fig. S2 for details.

In agreement with the hypothesis that at least one ribosome-bound nascent chain is required for the initial TF-PR dimerization reaction, longer constructs in which arrested TF-PR-RT nascent chains are more exposed outside the ribosome and hence better available for dimerization were more rapidly processed to mature ∼11 kDa PR, Supplemental Fig. S3a. Restoration of the PR/RT cleavage site (TLNF/PISG; same as in the L32 construct) made the arrested form of the L30 construct susceptible to autoproteolytic processing to mature PR, but with markedly slower kinetics than seen for the L32 construct, Supplemental Fig. S3b.

Rapid (2 min. labeling) pulse-chase analysis of the autoproteolysis reaction for the non-arrested L30[FL_c_] construct revealed a very different kinetic behavior, Fig. 5. In this case, the PR+linker+AP+tail cleavage product appeared together with the full-length chains already after a ∼2 min. chase, *i.e*., as soon as the 210-residue long full-length chains were produced (the translation rate in the PURE system is ≲ 1 aa/sec ^71,73^), and the relative amounts of cleaved *vs*. full-length products remained approximately constant during the chase (*c.f*., Fig. 3a, b). Thus, there is no detectable delay in the production of active PR for non-arrested constructs; nevertheless, there is only partial processing of the full-length chains. Again, this suggests that only ribosome-attached nascent chains are accessible to proteolysis, while released full-length chains are not.

**Figure 5.**
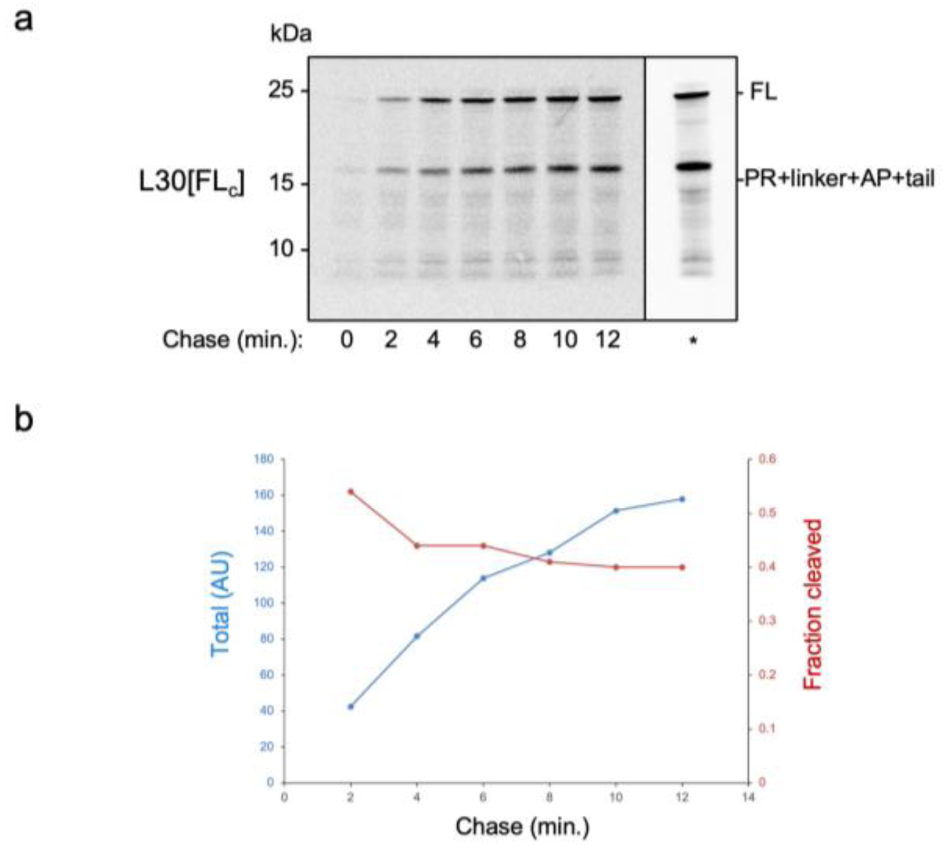
Rapid pulse-chase analysis of the L30[FL_c_] construct. (a) PURE *in vitro* translation reactions were carried out for 2 min. in the presence of [^35^S]-Met followed by a chase in the presence of excess non-radioactive Met. The lane indicated by * is for a 15 min. translation reaction in the presence of [^35^S]-Met with no chase, included for comparison. (b) Quantitation of the data in panel *a*, showing the total intensity of the 23 kDa + 16 kDa bands (arbitrary units, blue curve) and the fraction cleaved product (red curve).

To separate the effects of the initial, dimerization-dependent production of PR from the PR-mediated cleavage of nascent TF-PR-RT chains, we performed a ‘spiking’ experiment on L30[A_c_], L60(D^25^N)[A_c_], and L60(D^25^N)[FL_c_] by adding purified active PR (1 μM final concentration) to the translation reaction, either at the start of the 15 min. [^35^S]-Met labeling period, or together with the non-radioactive Met at the start of the chase period. For the ribosome-arrested L60(D^25^N) construct, Fig. 6a (lanes 1-8), inclusion of PR during the labeling period led to a reduction in both the *FL* and *A* products, with a concomitant appearance of both partial (PR+linker+AP+tail, TF+PR) and full (PR) cleavage products (compare lanes 3 and 6); a similar cleavage profile was seen when active PR was added at the start of the chase period, except that the *FL* product was now largely protease-resistant (compare lanes 4 and 7) and did not decrease in intensity during the 45 min. chase (compare lanes 7 and 8). Similar results, although with overall lower levels of proteolysis, were seen for the non-arrested L60(D^25^N)[FL_c_] construct (lanes 9-16): appearance of partial and full proteolysis products at early time points (lanes 14, 15), but no further processing during the 45 min. chase (lane 16), again demonstrating that both the *FL* and partial proteolysis products are largely refractory to proteolysis after they have been released from the ribosome. The 9.5 kDa band noted above appears even when no DNA was added to the PURE reaction (lanes 1, 2 and 9, 10), hence this band originates from the PURExpress system itself. For L30[A_c_] – which produces only marginal amounts of cleaved product even during a 45 min. chase, Fig. 3b – ‘spiking’ with purified PR led to a rapid appearance of the PR+linker+AP product, Fig. 6b, amounting to ∼50% of the total. Thus, ribosome-bound L30[A_c_] is a good substrate for PR but is very inefficient in forming the initial TF-PR dimers.

**Figure 6.**
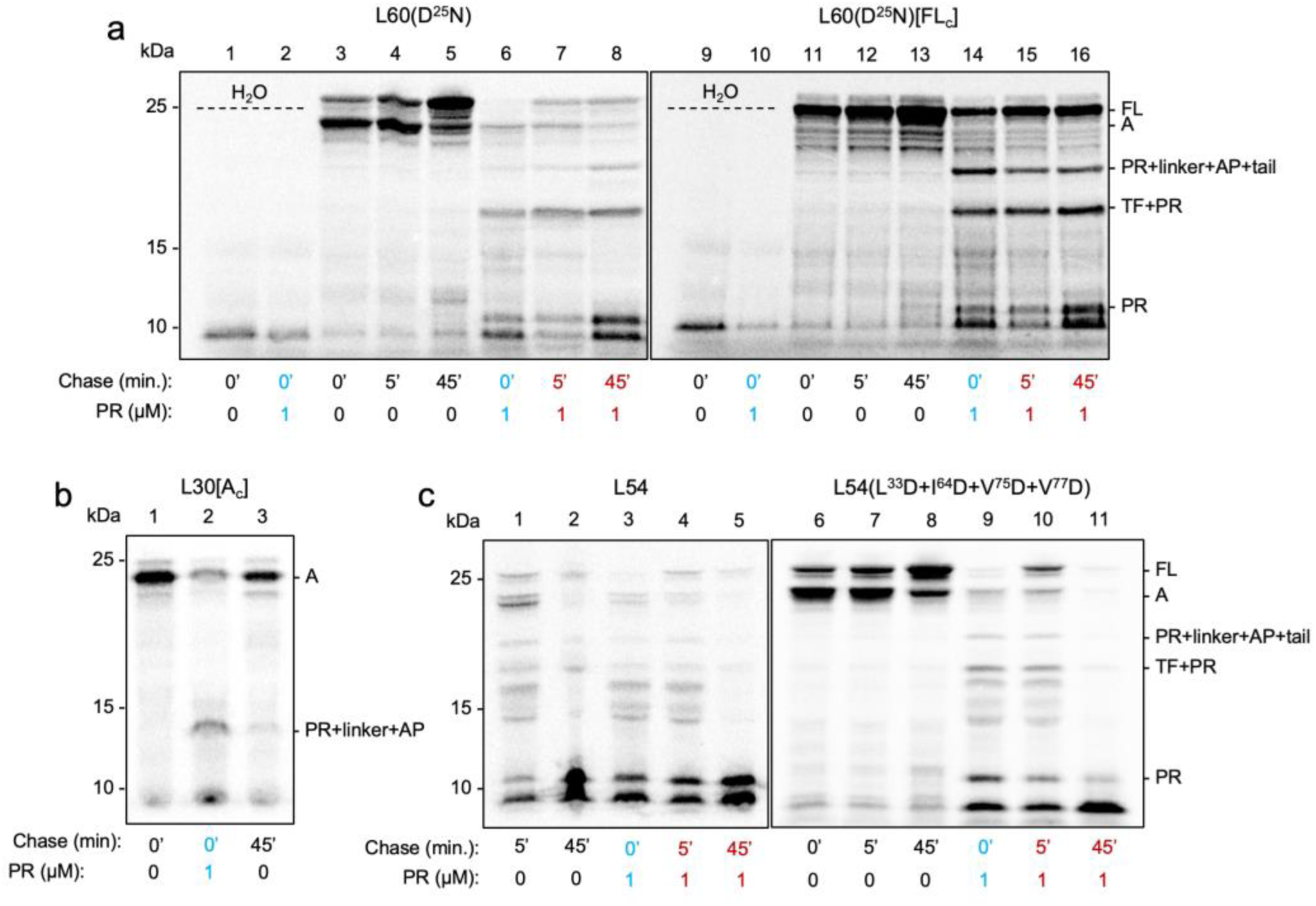
“Spiking” of the in vitro translation reaction with active PR. (a) Lanes 1-8: Inactive L60(D^25^N) was translated *in vitro* for 15 min. in the presence of [^35^S]-Met, followed by a chase in the presence of an excess of non-radioactive Met. Reactions were carried out in the presence or absence of purified, active PR at 1 μM final concentration, as indicated. Cyan labels indicate that PR was added at the beginning of the reaction, red labels indicate that PR was added at the beginning of the chase. The first two lanes, marked *H_2_O,* were reactions carried out without added construct DNA. Lanes 9-16: Same as in lanes 1-8, but for L60(D^25^N)[FL_c_]. (b) Same as in panel *a*, but for L30[A_c_]. (c) Same as in panel *a*, but for L54 (lanes 1-5) and L54(L^33^D+I^64^D+V^75^D+V^77^D) (lanes 6-11).

We also tested L54 and the presumably non-dimerizing, inactive mutant L54(L^33^D+I^64^D+V^75^D+V^77^D) in the spiking assay, Fig. 6c. L54 behaves as expected, *i.e*., both the *FL* and *A* products are largely proteolyzed to the 11 kDa product when PR is present during the labelling period (lane 3), whereas the *FL* product remains largely PR-resistant when PR is added at the start of the chase period (compare lanes 1 and 4, and lanes 2 and 6). In contrast, for L54(L^33^D+I^64^D+V^75^D+V^77^D), the *FL* product is sensitive to PR during the chase, and little *FL* remains after 45’ chase in the presence of PR (compare lanes 8 and 11). Thus, released *FL* product is largely PR-resistant in the dimerizing L54 construct, but not in the L54(L^33^D+I^64^D+V^75^D+V^77^D) mutant, suggesting that dimeric but not monomeric TF-PR-RT is resistant to cleavage by PR.

### The lag-time midpoint increases with the strength of the AP

In a final set of experiments, we tested the effects of the AP itself on the autoproteolytic reaction, comparing the SecM(*Ec*) AP to the much stronger SecM(*Ec*)-3W AP ^55^. As expected from the extent of translational pausing caused by the SecM(*Ec*) AP at low pulling forces ^54^, a slow conversion of the arrested to the full-length form (release-time midpoint 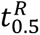 = 35 min) was seen for the catalytically inactive L60(D^25^N) mutant, Fig. 7a (also evident in Fig. 6a). Using the stronger SecM(*Ec*)-3W AP that retains the nascent chain on the ribosome more efficiently than does the wildtype SecM(*Ec*) AP, the conversion of the arrested to the full-length form of L60(D^25^N)-3W took even longer (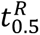 = 61 min), Fig. 7b.

**Figure 7.**
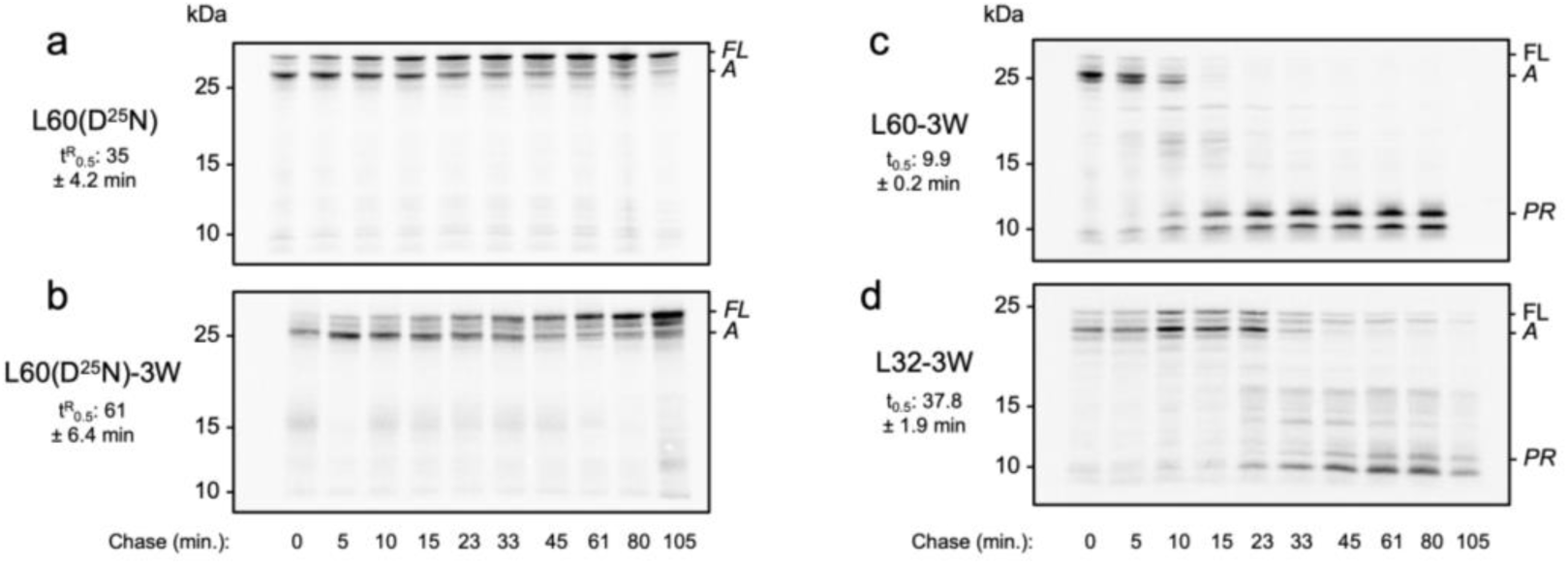
Pulse-chase analysis of SecM(Ec)-3W AP constructs. Pulse chase experiments were performed as in Fig. 4. *FL*, *A*, and PR products are indicated. *3W* indicates that the strong SecM(*Ec*)-3W mutant AP was used. Release-time midpoints (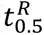) are indicated in panels *a* and *b*, and lag-time midpoints in panels *c* and *d*. (see Supplemental Fig. S2 for quantitations).

With the stronger SecM(*Ec*)-3W AP, the long L60 construct in which the entire PR domain is exposed outside the ribosome in the arrested nascent chain still underwent rapid autoproteolytic processing but with a two-fold increase in the lag-time midpoint (*t*_0.5_ = 9.9 min. *vs*. 5.3 min. with the SecM(*Ec*) AP), Fig. 7c, which parallels the ∼2-fold increase in the release-time midpoint between the SecM(*Ec*) and the SecM(*Ec*)-3W APs. Autoproteolysis of the short L32 construct in which the C-terminal end of the PR domain is sequestered inside the exit tunnel in the arrested nascent chain was also delayed by the SecM(*Ec*)-3W AP, again with a ∼2 fold increase in *t*_0.5_ from 17 min. for the SecM(*Ec*) AP to 38 min. for the SecM(*Ec*)-3W AP, Fig. 7d. Thus, the release-time midpoint appears to set the rate at which active PR is produced, implying that the initially formed TF-PR-RT dimers must be released from the ribosome in order to start cleaving other ribosome-bound TF-PR-RT nascent chains.

## Discussion

We have analyzed the cotranslational folding and proteolysis of frameshifted TF-PR-RT constructs of different lengths using both FPA and autoproteolysis assays in the *E. coli*-derived PURE *in vitro* transcription-translation system. In short, we find that the PR monomer starts to fold in the ribosome exit tunnel when the end of its C-terminal helix is ∼30 residues away from the PTC (Fig. 2b). Previous studies suggest that at this depth the exit tunnel can accommodate folding of protein domains of up to ∼45 amino acids long, whereas domains the size of the PR monomer (∼99 amino acids) can only fold when ≳ 35 residues away from the PTC ^74^. Hence, the observed folding event is unlikely to involve the whole monomer, and in fact is not affected when core hydrophobic residues in folded PR are mutated, Fig. 2b. However, mutations in the C-terminal helix reduce the amplitude of the peak in the FP at *L* = 30 residues (Fig. 2c), suggesting that a main contribution to the peak is from folding of the C-terminal helix.

A second peak of lower amplitude is seen at *L* = 52-56 residues, at *L* values too large to represent the folding of the PR monomer ^74^. At such large *L* values, the PR part of the TF-PR-RT nascent chain should be sufficiently far out of the ribosome to be able to dimerize with another TF-PR-RT molecule ^75^, either in *cis* (*i.e*., with TF-PR-RT being synthesized on the same mRNA by the trailing ribosome) or in *trans* (*i.e*., with TF-PR-RT being synthesized on a different mRNA, or with released full-length TF-PR-RT). Both kinds of cotranslational dimerization have been described before ^69,75^. As expected for a dimerization event, mutations in the hydrophobic core of PR reduce the amplitude of this peak (Fig. 2b).

We have also found that catalytically active PR dimers can form autoproteolytically from the TF-PR-RT protein in the PURE *in vitro* transcription/translation system, as has been shown before for the rabbit reticulocyte *in vitro* translation system ^31^. Unexpectedly, however, processing of the TF-PR-RT protein is seen only for arrested, ribosome-attached chains, while released full-length and partially processed chains remain uncleaved even after a 105 min. chase (Fig. 4). Similarly, as seen for the L60(D^25^N) and L60(D^25^N)[FL_c_] constructs, full-length and proteolysis products not bound to the ribosome are largely resistant to purified, active PR added to the translation reaction, except for the presumably non-dimerizing L54(L^33^D+I^64^D+V^75^D+V^77^D) mutant (Fig. 6). Thus, PR cleaves TF-PR-RT while it is being translated but not after release from the ribosome.

Our results further suggest that the initial dimerization of TF-PR-RT monomers involves at least one ribosome-bound nascent chain. Thus, L30[FL_c_] produces ample amounts of active PR already after a 2 min. translation reaction followed by a 2 min. chase (Fig. 5), while the closely related L30[A_c_] produces only trace amounts of active PR even after a 15 min. translation reaction followed by a 45 min. chase (Fig. 3b). The only difference between these two constructs is the 23-residue C-terminal tail present in L30[FL_c_], which causes the TF-PR portion to be exposed outside the ribosome exit tunnel for ∼10-15 sec before chain termination in L30[FL_c_], but not in L30[A_c_].

In contrast to FL_c_ and A_c_ controls, arrested nascent chains spend a considerable time in the ribosome-bound state before being released, either by slow release of the AP or by autoproteolytic cleavage at the PR/RT cleavage site. As seen by comparing the initial lag-time midpoint in the onset of autocatalytic processing (*t*_0.5_) between constructs arrested by the medium-strong SecM(*Ec*) AP and the very strong SecM(*Ec*)-3W AP (Fig. 7), *t*_0.5_ appears to be set by the release-time midpoint (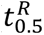), again implying that the rate-limiting step in the production of active PR for constructs with an active AP is the release of preformed TR-PR-RT dimers from the ribosome.

Thus, as illustrated in Fig. 8a, under the *in vitro* conditions employed here, the TF-PR-RT protein can, to a first approximation, engage in the initial dimerization reaction required to produce active PR, and is susceptible to PR-mediated cleavage, only when still attached to the ribosome. Released full-length or partially processed chains either aggregate or, more likely, form low-activity TF-PR-RT dimers that are not in themselves substrates for PR, and hence remain unprocessed. This model does not require the initial TF-PR-RT dimer to self-cleave ^41–45^ in order for the first PR molecules to be released, because monomeric, ribosome-bound nascent TF-PR-RT substrates will be available on neighboring ribosomes.

**Figure 8.**
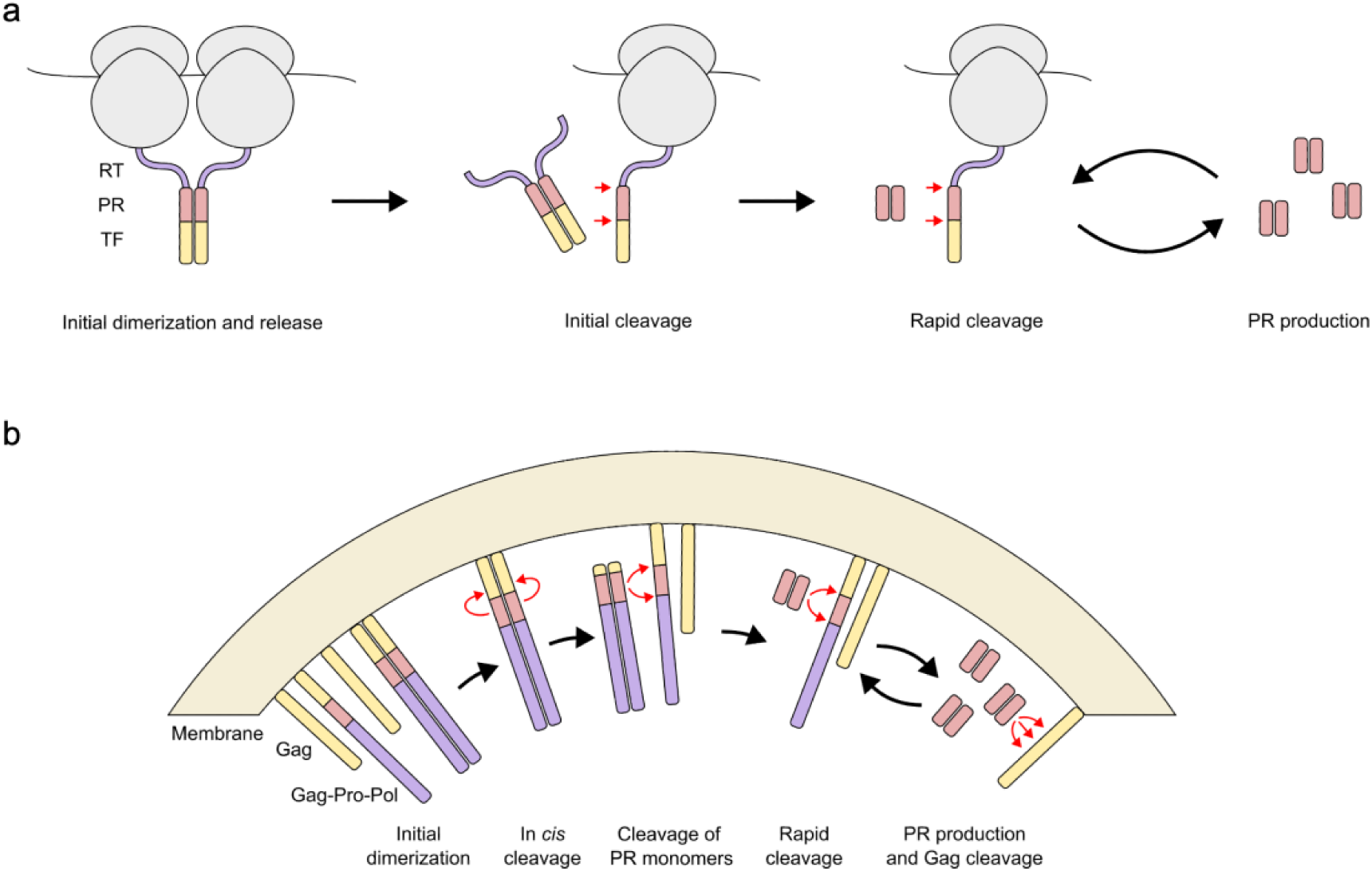
Models for autocatalytic processing of TF-PR-RT constructs. (a) Cotranslational processing during *in vitro* translation (this work). (b) Protease activation during virion assembly.

Translating these findings to the situation in the HIV-1 infected cell, the low frequency (-5%) of the -1 frameshifting required to produce Gag-Pro-Pol appears critical, as only 1 in ∼400 ribosomes would both carry nascent Gag-Pro-Pol and have a neighboring ribosome also carrying Gag-Pro-Pol and thus be able to release preformed Gag-Pro-Pol dimers upon chain termination; in addition, such dimers would be released far away from other ribosomes carrying monomeric TF-PR-RT nascent chains, which would further reduce the rate of the initial cleavage reaction. Compared to the frameshifted TF-PR-RT constructs studied here, the initial lag time in the onset of autocatalytic processing (*t*_0.5_) of wild-type Gag and Gag-Pro-Pol in a newly infected cell would therefore be up to two to three orders of magnitude longer than what is observed here, preventing the appearance of active PR dimers in the cytosol.

While this experimental system involves an *in vitro* translation setting, there are elements of its behavior that suggest a model for protease activation during virion assembly. There are approximately 2,500 Gag molecules and thus 125 Gag-Pro-Pol molecules in a budding virion anchored on the inner face of the viral membrane ^2^. If Gag-Pro-Pol molecules enter the budding particle as monomers (or as low complexity oligomers with Gag), then the chance juxtaposition of two Gag-Pro-Pol molecules during assembly, possibly potentiated by the curvature of budding, would allow an infrequent dimerization event (at least one per virion on average). Such a dimerization event would be followed by upstream cleavage at SP2/NC and in TF in *cis* ^31^ to release a TF-PR-RT-IN precursor dimer from the viral membrane. This dimeric form of the precursor could then diffuse within the lumen of the virion to allow interaction with or cleavage of monomeric Gag-Pro-Pol that would then generate PR monomers. The PR monomers would form the highly active dimers needed to carry out all of the cleavage events needed for virion maturation, Fig. 8b.

## Acknowledgments

We thank Dr. Celia Schiffer (University of Massachusetts Chan Medical School) for generously providing purified HIV-1 protease, Ean Spielvogel for assistance with oligonucleotide design, and Dr. Rickard Hedman (Stockholm University) for programming and maintenance of the EasyQuant software. This work was supported by grants from the Knut and Alice Wallenberg Foundation (2017.0323), the Novo Nordisk Fund (NNF18OC0032828), and the Swedish Research Council (621-2014-3713) to GvH, and by NIH award R01 GM135919 to RS (Celia Schiffer, PI).

## Data availability statement

Python scripts used for fitting pulse-chase curves are available upon reasonable request.

---

**Supplemental Figure S1.**
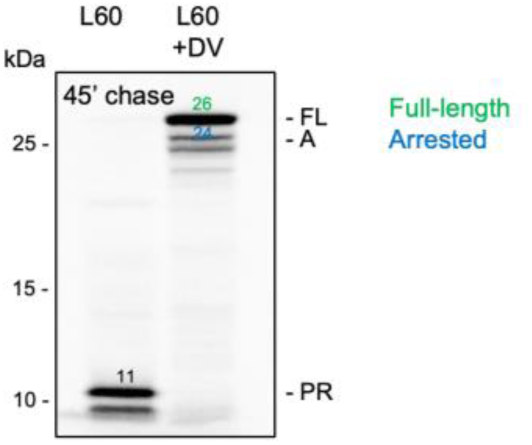
Effect of darunavir (DV) on the autoproteolysis of the L60 construct. L60 was translated *in vitro* in the presence of [^35^S]-Met for 15 min. and then chased in the presence of excess non-radioactive Met for 45 min, in the absence and presence (+DV) of 850 μM darunavir. Equal concentrations of DMSO were included in the ±DV reactions. Mw’s are indicated, color-coded as in Fig. 2a.

**Supplemental Figure S2.**
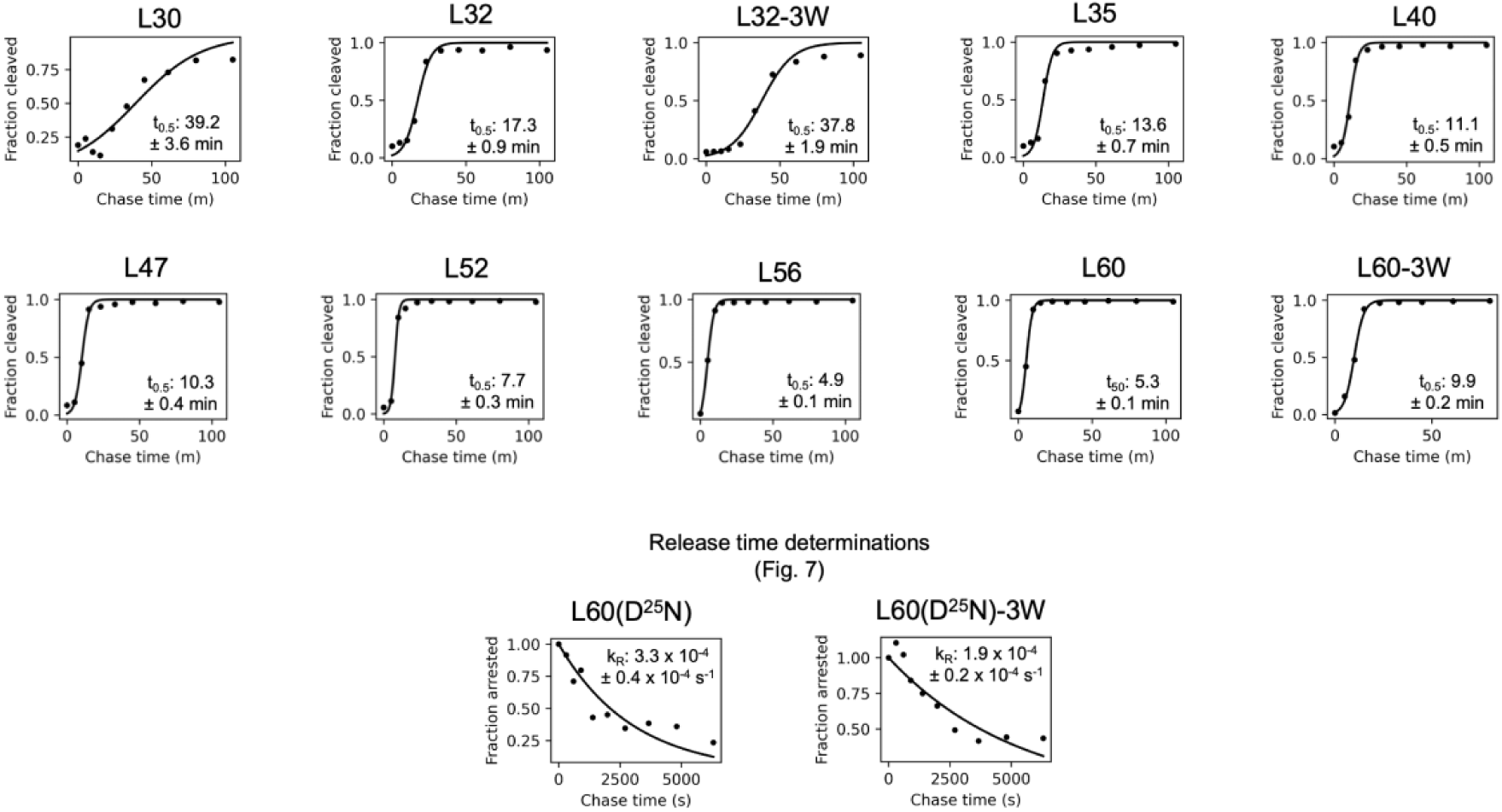
Determination of lag-time (t_0.5_) and release-time (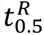) midpoints. The intensities of bands representing the arrested fraction and the mature PR in Fig. 4 were fit to a two-state sigmoidal equation to determine *t*_0.5_values and standard errors. For D^25^N constructs, plots and fits used to determine 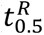 values are shown ± the standard error of the fitted value. See Methods for details.

**Supplemental Figure S3.**
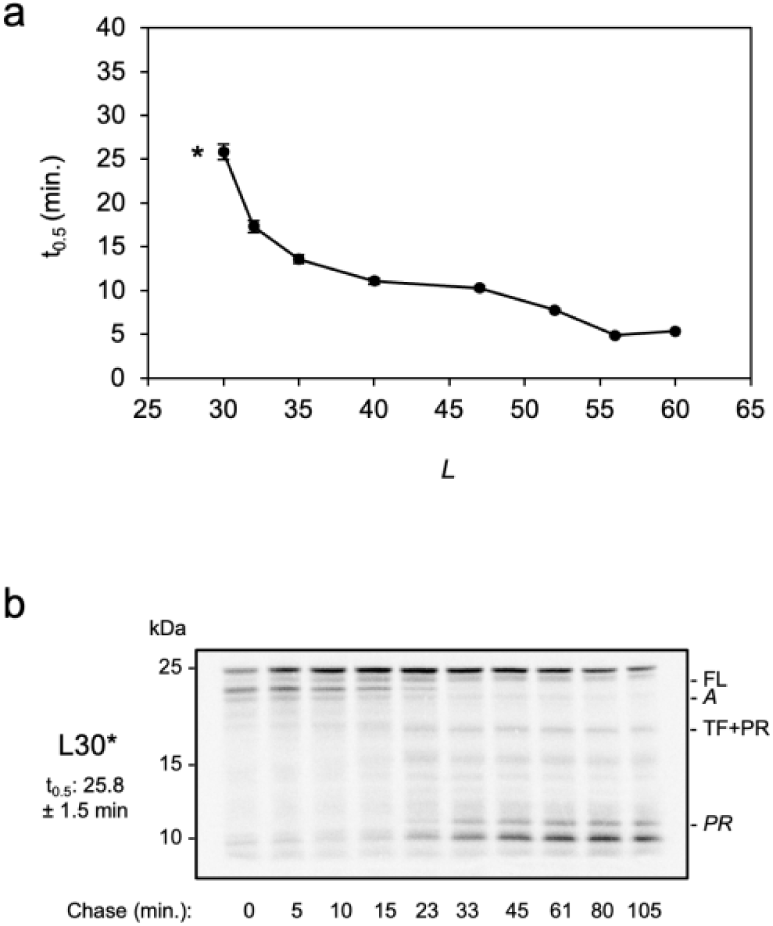
Lag-time midpoints vary with L. (a) *t*_0.5_ values for the constructs shown in Fig. 4. The value for *L* = 30 residues (indicated by *) is for the construct shown in panel *b*. (b) Pulse-chase analysis (15 min. pulse) of the mutated L30* construct with an active PR/RT cleavage site (TLNF/PISG).

